# Single-nucleus transcriptome analyses uncover a dynamic transcriptional landscape of soybean roots in response to soybean cyst nematode infection

**DOI:** 10.1101/2025.08.06.668855

**Authors:** Pan-Pan Bai, Xin-Ye Jia, Luying Chen, Yatao Han, Shaojie Han, Lin Weng, Xianzhong Feng

**Affiliations:** Institute of Virology and Biotechnology, Zhejiang Academy of Agricultural Sciences, Hangzhou 310021, Zhejiang, China; Research Center for Life Sciences Computing, Zhejiang Laboratory, Hangzhou 311100, China; State Key Laboratory of Rice Biology and Breeding, Key Laboratory of Biology of Crop Pathogens and Insects of Zhejiang Province, Institute of Biotechnology, Zhejiang University, Hangzhou 310058, Zhejiang, China; Key Laboratory of Soybean Molecular Design Breeding, National Key Laboratory of Black Soils Conservation and Utilization, Northeast Institute of Geography and Agroecology, Chinese Academy of Sciences, Changchun 130102, China

**Keywords:** soybean, soybean cyst nematode, syncytium, single-nuclei RNA sequencing, transcriptomic atlas, nematode resistance

## Abstract

Soybean cyst nematode (SCN, *Heterodera glycines*) causes substantial yield losses by inducing the formation of specialized feeding structure called syncytia, which arise from extensive reprogramming of host root cells. Despite their pivotal role in SCN parasitism, the molecular mechanisms governing syncytium formation and function remain poorly understood, largely due to the technique challenges in precisely isolating these rare cell types. Here, we employed single-nucleus RNA sequencing (snRNA-seq) to generate a comprehensive, high-resolution transcriptomic atlas of SCN-infected soybean roots, comparing two resistant cultivars (PI 88788 and Forrest) with a susceptible cultivar (Williams 82) across multiple infection stages. By leveraging the well-characterized syncytium marker *GmSNAP18*, we successfully identified and transcriptionally profiled syncytial cells, and validate the spatial expression of known SCN-resistant genes. Our analyses revealed strong induction of resistance-associated genes specifically enriched in syncytia from resistant cultivars. Functional validation further demonstrated that overexpression of three novel candidate genes significantly suppressed SCN development. Pseudotime trajectory analyses traced syncytium development back to procambial cell origins, uncovering distinct, cultivar-specific developmental pathways. Collectively, this work provides a foundational single cell transcriptomic resource, advances our understanding of syncytial biology and SCN resistance mechanisms, and identifies promising molecular targets for engineering nematode-resistant soybean varieties.

## Introduction

Soybean (*Glycine max* [L.] Merr.) is a globally important legume crop, providing essential plant-based protein and oil for both human consumption and animal feed. However, soybean production is severely threatened by the soybean cyst nematode (SCN, *Heterodera glycines*), which causes significant yield losses worldwide with annual economic damage in China alone estimated at $120 million (Bradley et al., 2021; Koenning and Wrather, 2010; Li et al., 2011). The critical step in SCN parasitism is the formation of syncytia, specialized feeding structures derived from the reprogramming of host root cells (Acedo et al., 1984; Mitchum, 2016). Despite recent advances, the gene expression profiles within syncytial cells and the molecular mechanisms underlying their development remain poorly characterized.

Syncytia originate from an initial syncytial cell (ISC), typically located within the vascular cylinder and derived from procambial or pericycle cells (Anjam et al., 2020; Kloc and Uosef, 2024). These cells develop into multinucleate feeding structures via nematode-induced protoplast fusion and localized cell wall dissolution. As the development of syncytial cells, they are characterized by thickened cell walls, dense cytoplasm, and elevated ribosomal and endoplasmic reticulum activity, functioning as strong metabolic sinks that support nematode growth (Gipson et al., 1971; Jones, 1981; Sobczak et al., 1997b). Although SCN is capable of initiating syncytium formation in resistant soybean cultivars, these structures typically undergo necrosis shortly after their establishment.

Previous studies using laser microdissection have identified syncytium-specific genes involved in cell wall remodeling, hormone signaling, and nutrient transport (Liu and Mitchum, 2025). Comparative transcriptomic analyses between resistant and susceptible cultivars, as well as between syncytia and whole infected roots, have revealed key expression differences (Goode and Mitchum; Ithal et al., 2007; Klink et al., 2007a; Szakasits et al., 2009). However, the lack of single-cell resolution has limited our ability to distinguish transcriptional profiles of syncytia from adjacent root cells, thereby constraining our understanding of the regulatory mechanisms governing syncytial development. Recent advances in snRNA-seq have enabled the construction of a high-resolution transcriptomic atlas and gene regulatory networks in diverse plant tissues (Guo et al., 2025; Lee et al., 2023; Liu et al., 2023; Zhang et al., 2025). For instance, integration of snRNA-seq with spatial transcriptomics revealed vein-specific immune responses in rice leaves to *Magnaporthe oryzae*, including activation of the resistance gene *OsHKT9* (Wang et al., 2025). These advances highlight the potential of snRNA-seq to elucidate the cellular and molecular basis of syncytial development during SCN infection.

Soybean SCN resistance is a complex quantitative trait, with more than 300 QTLs cataloged in SoyBase (https://www.soybase.org/). Nevertheless, among the identified loci conferring resistance, resistant to *Heterodera glycine* 1 (*Rhg1*, located on chromosome 18) and *Rhg4* (located on chromosome 8) have been most extensively characterized to date. The *Rhg1* locus contains three genes within a 31-kb genomic segment: *Glyma.18G022400*, encoding a putative amino acid transporter (*GmAAT*); *Glyma.18G022500* (*GmSNAP18*), encoding an α-soluble NSF attachment protein (α-SNAP), which is involved in vesicle trafficking; and *Glyma.18G022700*, encoding a WIP protein with a WI12 damage-induced domain. Resistance conferred by *Rhg1* is associated with copy number variation of these genes (Cook et al., 2012). The *Rhg4* locus encodes *Glyma.08g108900* (*GmSHMT08*), a serine hydroxymethyltransferase identified from the Peking cultivar (Liu et al., 2012). More recently, *GmPR10-09g* (*Glyma.09G040400*), a member of PR10 protein family, has also been implicated in SCN resistance (Deng et al., 2024). Other studies have revealed that SCN resistance is also influenced by genes involved in exocytosis, phytohormone signaling, and miRNA-mediated regulatory pathways (Feng et al., 2022; Lin et al., 2013; Noon et al., 2019). Notably, *GmSNAP18* and *GmPR10-09g* exhibit syncytium-specific expression patterns, supporting the hypothesis that spatially restricted genes in resistant cultivars plays a central role in regulating soybean-SCN interactions (Bayless et al., 2016; Bayless et al., 2019; Deng *et al*., 2024).

To gain single-cell resolution insights into the molecular dynamics of SCN infection, we selected two classical SCN-resistant cultivars, PI 88788 (carrying high-copy *rhg1-b*) and Forrest (carrying low-copy *rhg1-a* and *Rhg4*), along with a susceptible cultivar, Williams 82 (harboring single-copy *rhg1-c*). Root segments encompassing the SCN infection zone were collected at 1, 3, and 7 days post inoculation (dpi), corresponding to key stages of parasitism and syncytium development, and subjected to snRNA-seq. Using this approach, we aimed to construct a dynamic, high-resolution single-nucleus transcriptomic atlas that captures spatial and temporal gene expression patterns during SCN infection. We identified some novel genes specifically enriched in syncytial cells of resistant cultivars and conducted functional assays to validate their roles in mediating resistance. In addition, we reconstructed syncytium developmental trajectories, revealing key regulatory genes involved in their initiation and differentiation. Collectively, our study provides novel insights into the molecular mechanisms underlying SCN resistance and syncytial development, offering a valuable resource for understanding plant-nematode interactions at the single-cell resolution and facilitating the genetic improvement of soybean for nematode resistance.

## Results

### Construction of a single-nucleus transcriptomic atlas of soybean roots during SCN infection

To comprehensively capture the molecular responses of soybean roots to SCN infection, we performed snRNA-seq on root infection zones collected from three cultivars, two resistant (PI 88788 and Forrest) and one susceptible (Williams 82) at 1, 3, and 7 dpi. To ensure reproducibility, two replicates were prepared for each sample (Figure 1A). Following stringent quality control, we obtained a total of 172,199 high-quality nuclei, with a median of 1,870 expressed genes and 2,732 unique molecular identifiers (UMIs) per nucleus (Table S1; Figure S1A, B).

**Figure 1.**
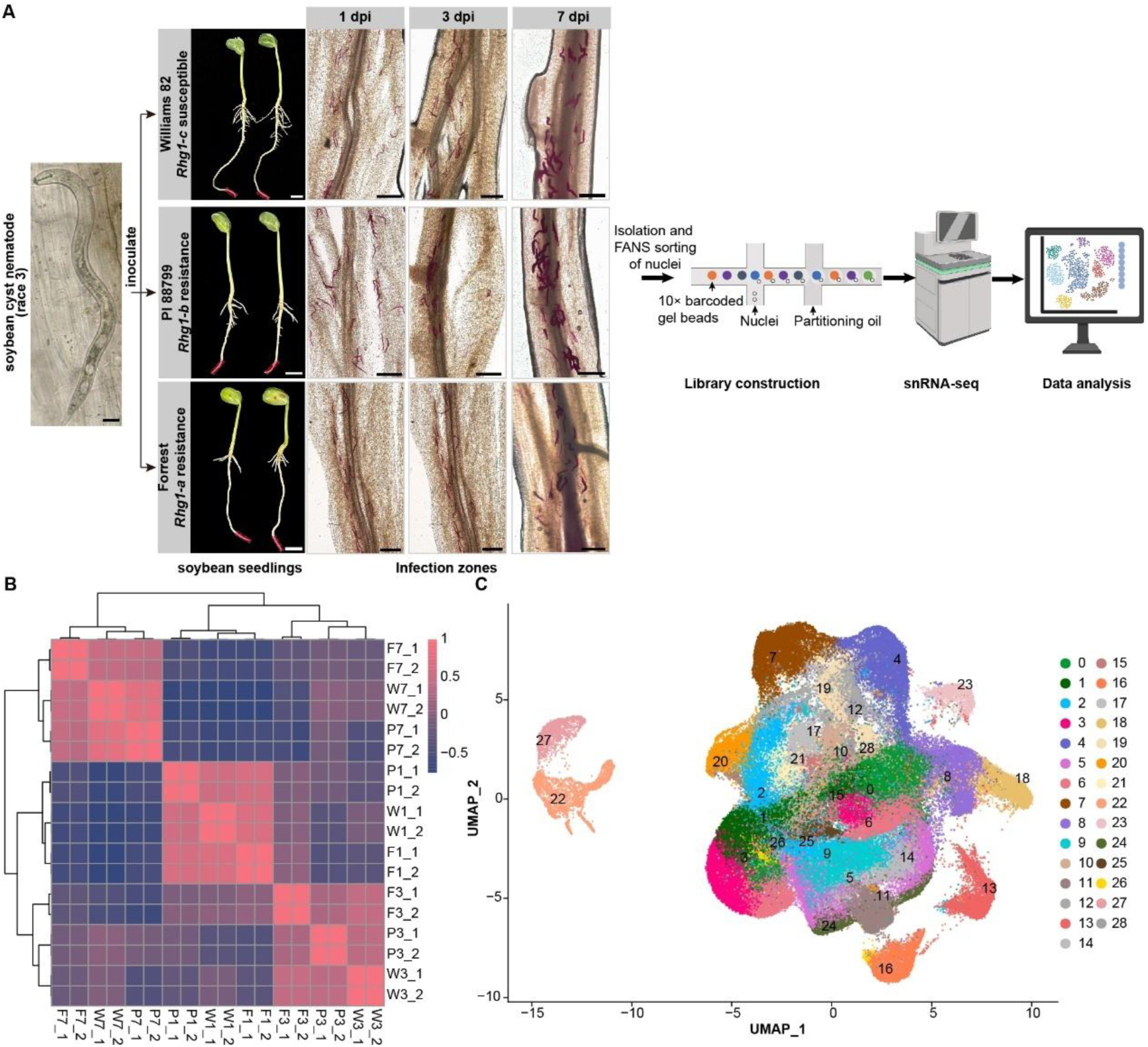
The transcriptional atlas of soybean roots after inoculation by soybean cyst nematode. (A) Workflow diagram illustrating the snRNA-seq workflow. Approximately 400 infective J2 SCN (bar = 20 µm) were applied 1 cm above the soybean seedling root tip (bar = 1 cm). The infection zones (bar = 500 µm) of Williams 82, PI 88788, and Forrest at 1, 3, and 7 days post inoculation (dpi) were collected to perform single-nucleus RNA sequencing, respectively. (B) Heatmap displaying Pearson correlation coefficients among samples. (C) UMAP visualization of 29 clusters from soybean roots and SCN. F1: Forrest at 1 dpi; F3: Forrest at 3 dpi; F7: Forrest at 7 dpi; P1: PI 88788 at 1 dpi; P3: PI 88788 at 3 dpi; P7: PI 88788 at 7 dpi; W1: Williams 82 at 1 dpi; W3: Williams 82 at 3 dpi; W7: Williams 82 at 7 dpi.

Nuclei were classified based on gene content: those with >80% soybean-derived transcripts were defined as soybean-origin. In total, we identified 168,117 soybean-origin nuclei and 4,082 SCN-origin nuclei (Table S1; Figure S1C). Biological replicates exhibited high reproducibility, with Pearson correlation coefficients (PCCs) ranging from 0.97 to 1.0. Interestingly, gene expression profiles were more strongly correlated across time points than between cultivars, suggesting a largely conserved transcriptional response to SCN infection (Figure 1B).

Integration of all 18 snRNA-seq datasets revealed 29 transcriptionally distinct cell clusters, visualized using uniform manifold approximation and projection (UMAP) (Figure 1C). The consistent distribution of clusters across cultivars indicates that SCN infection induced similar cellular composition and transcriptional responses, regardless of resistance genotype (Figure S2).

### Comprehensive cell type annotation of soybean root clusters

To accurately annotate root cell types, we employed four complementary strategies. First, we performed manual annotation based on known cell type-specific markers from *Glycine max* and *Arabidopsis thaliana*, which assigned identities to all clusters except cluster 2 (Table S2, Table S3) (Bayless *et al*., 2016; Bayless *et al*., 2019; Cervantes-Pérez et al., 2024; Denyer et al., 2019; Jean-Baptiste et al., 2019; Ryu et al., 2019; Shahan et al., 2022; Sun et al., 2023; Zhang et al., 2019). Cluster 17 was annotated as syncytium based on strong expression of *GmSNAP18*, a well-characterized *Rhg1* locus gene specifically expressed in syncytial feeding cells(Bayless *et al*., 2016; Bayless *et al*., 2019; Han et al., 2023; He et al., 2025; Lakhssassi et al., 2020) (Figure 2A). Second, we used the “Cell Type Predictor” in the scPlantDB database (He et al., 2024) to calculated similarity scores between the *Arabidopsis* orthologs of highly differentially expressed genes in each cluster and *Arabidopsis* root cell type markers. The highest similarity scores in the mature root zone were used to assign predicted identities (Figure 2B; Table S4). Third, 11,158 anchors were found between our dataset and a recently released soybean root single-cell atlas using Seurat (Hao et al., 2024; Zhang *et al*., 2025), and they were used to classify the query cell based on reference data (Figure 2C). Fourth, we applied the marker gene-based classifier Garnett (Pliner et al., 2019), to predict the cell types based on the highly expressed cell-type-specific marker genes in soybean and *Arabidopsis* (Figure 2D).

**Figure 2.**
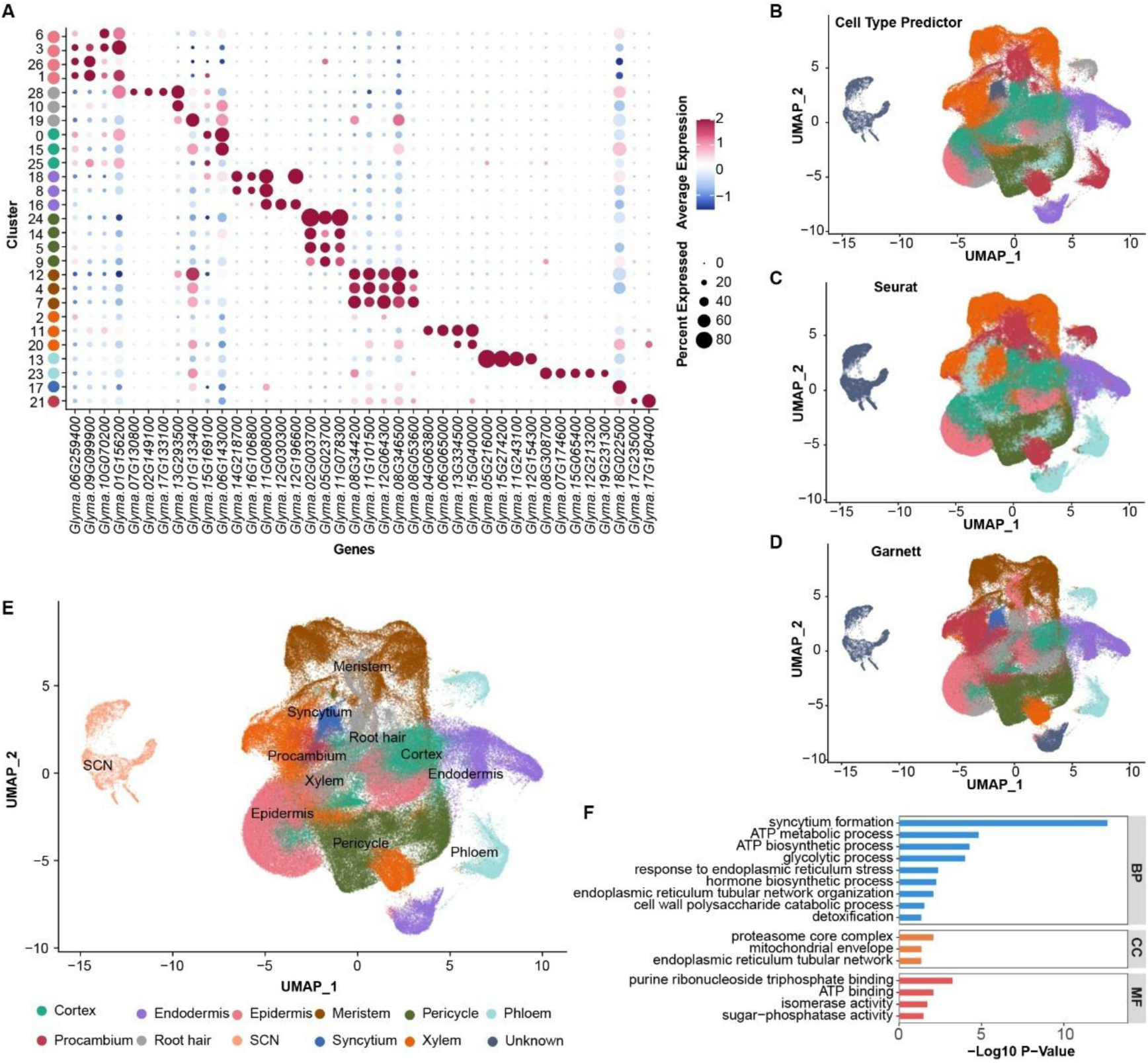
11 cell types in soybean and soybean cyst nematode were annotated using four complementary approaches. (A) Marker gene expression patterns for annotated clusters based on known soybean and *Arabidopsis thaliana* cell-type-specific genes. (B) UMAP visualization of cell types predicted using the “cell type Predictor” tool in the scPlantDB. (C) UMAP visualization of cell types predicted using Seurat. (D) UMAP visualization of cell types predicted using Garnett. (E) UMAP visualization of 11 cell types annotated in soybean roots and soybean cyst nematode. (F) Enriched GO terms of syncytium-specific genes.

Consistent results across all methods validate the accuracy of our annotations (Figure 2A–D; Table S4). In total, we identified 10 primary root cell types: epidermis (clusters 1, 3, 6, 26), root hair (10, 19, 28), cortex (0, 15, 25), endodermis (8, 16, 18), meristem (4, 7, 12), pericycle (5, 9, 14, 24), procambium (21), syncytium (17), phloem (13, 23), xylem (2, 11, 20), plus two nematode-derived clusters (22, 27) (Figure 2E; Table S4). Traditional techniques such as laser microdissection have limited capacity to isolate syncytia, thereby constraining identification of SCN-responsive genes (Ithal *et al*., 2007; Klink *et al*., 2007a; b; Szakasits *et al*., 2009). Our snRNA-seq approach overcame this limitation, captured 4,141 syncytial cells and identified 1,378 putative syncytium-specific genes, including well-known genes such as *GmSNAP18* and *GmPR10-09g*. Functional enrichment analysis revealed these genes were associated with ATP biosynthesis, glycerol metabolism, detoxification, and hormone biosynthesis (Figure 2F), underscoring the elevated metabolic activity and nutrient demands of syncytial cells during nematode parasitism.

### Spatial distribution of known SCN resistance genes across root cell types

To validate cell type annotations and explore the spatial expression of known SCN resistance loci, we examined the transcriptional profiles of key resistance genes across annotated root cell types. Among the *Rhg1* associated genes, *GmSNAP18* (*Glyma.18G022500*) was strongly enriched in the syncytium of PI 88788 (Figure 3A), consistent with previous *in situ* hybridization results (Lakhssassi *et al*., 2020). This gene has also been shown to hyperaccumulate in the syncytia of Fayette, a PI 88788-derived resistant cultivar (Bayless *et al*., 2016). Two additional *Rhg1* genes, *GmAAT* (*Glyma.18G022400*) and *GmWI12* (*Glyma.18G022700*), exhibited preferential expression in the epidermis and endodermis of PI 88788, respectively (Figure 3B, C). The epidermal enrichment of *GmAAT* is consistent with its previously reported accumulation in SCN-penetrated cells (Han *et al*., 2023), implicating its response in early infection. At the *Rhg4* locus, *GmSHMT08* (*Glyma.08G108900*), a gene conferring Peking-type resistance, showed broad expression across root tissues, with notable enrichment in the xylem of PI 88788 and Forrest, and increased expression in the syncytium of Forrest at 1 dpi (Figure 3D).

**Figure 3.**
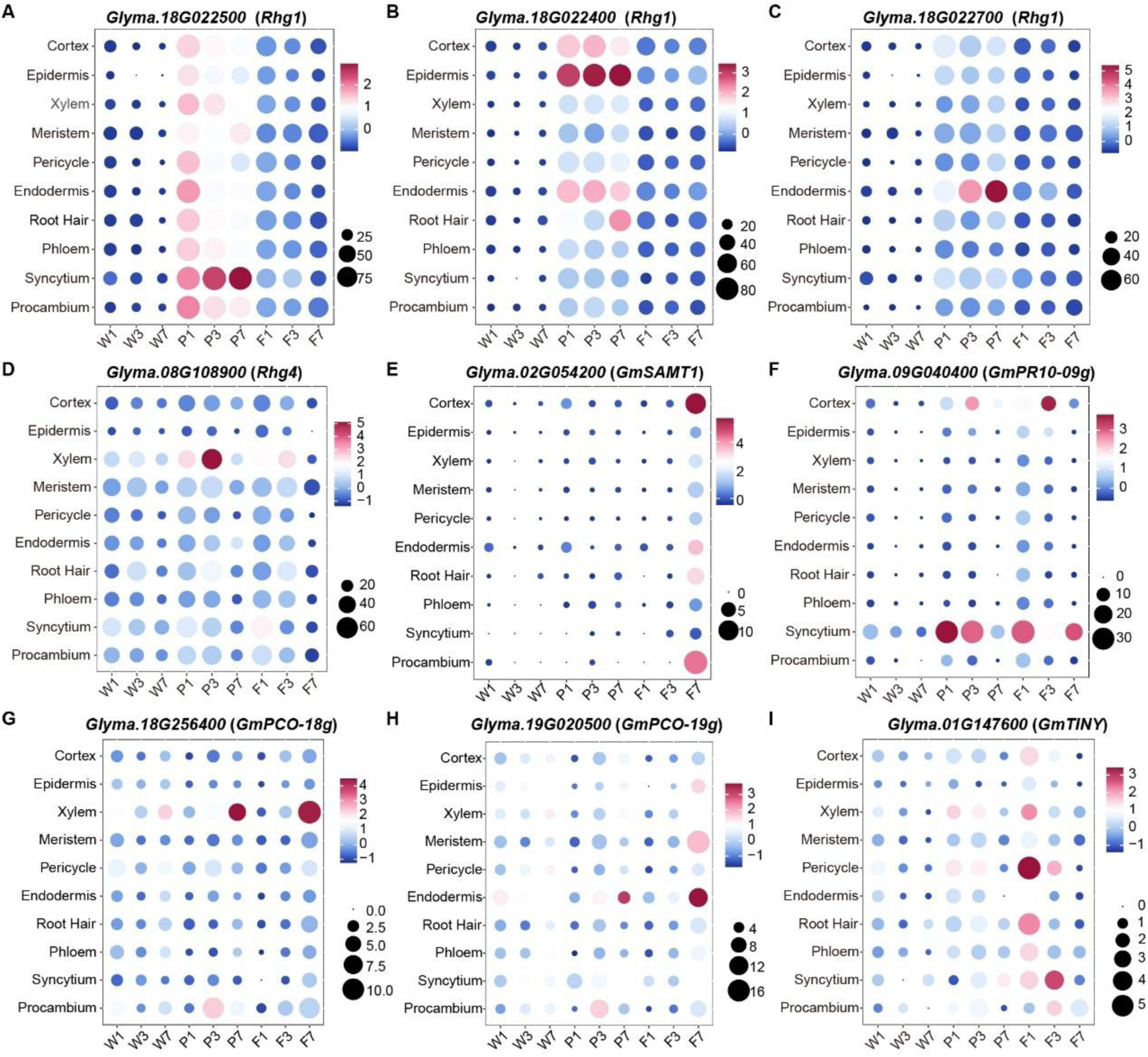
Known SCN resistance genes exhibited diverse root expression patterns. (A) –(C). Dot plots showing expression patterns of the functional genes at the *Rhg1* in each sample. (D) Dot plot showing expression pattern of the functional gene at the *Rhg4* in each sample. (E) Dot plot showing expression pattern of *Glyma.02G054200* in each sample. (F) –(H) Dot plot showing expression pattern of functional genes reported by Deng *et al*., 2024 in each sample. (I) Dot plot showing expression pattern of *Glyma.01G147600* in each sample. The color of dots indicates the average level of each gene, and the size of the dots represents the expression proportion of genes in each cell type. F1: Forrest at 1 day post inoculation (dpi); F3: Forrest at 3 dpi; F7: Forrest at 7 dpi; P1: PI 88788 at 1 dpi; P3: PI 88788 at 3 dpi; P7: PI 88788 at 7 dpi; W1: Williams 82 at 1 dpi; W3: Williams 82 at 3 dpi; W7: Williams 82 at 7 dpi.

The *salicylic acid methyltransferase 1* (*GmSAMT1*, *Glyma.02G054200*) (Lin et al., 2016; Lin *et al*., 2013) peaked in the cortex of Forrest at 7 dpi (Figure 3E), while *GmPR10-09g* (*Glyma.09G040400*) showed elevated expression in syncytia of resistant cultivars and in the cortex at 3 dpi (Figure 3F). Interestingly, two plant cysteine oxidase (*PCO*) genes (*Glyma.18G256400*, *GmPCO-18g*; *Glyma.19G020500*, *GmPCO-19g*) show significant upregulation in the xylem and endodermis in two resistant cultivars at 7 dpi, rather than high expression at the nematode feeding site as previously described (Figure 3G, H) (Deng *et al*., 2024). Additionally, the AP2/ERF transcription factor *GmTINY* (*Glyma.01G147600*) was significantly enriched in pericycle at 1 dpi and in the syncytium at 3 dpi in Forrest (Figure 3I) (He *et al*., 2025). Soybean *CLE* receptors, which have been linked to nematode resistance (Guo et al., 2015), displayed diverse expression patterns across root tissues (Figure S3). Collectively, these results demonstrate that key SCN resistance genes exhibit spatially distinct expression across multiple root cell types, including the epidermis, endodermis, cortex, and vascular system, in addition to the syncytium. This spatial complexity suggests that SCN resistance involves coordinated, multi-cellular defense networks rather than being confined to the nematode feeding site.

### Identification of novel genes in syncytial cells responsive to SCN

Syncytial transcriptional profiles diverge markedly between resistant and susceptible soybean-SCN interactions (Klink *et al*., 2007a). To identify genes contributing to resistance, we performed differential expression analysis on syncytial cells from the SCN-susceptible Williams 82 and the resistant cultivars PI 88788 and Forrest, across matched infection time points. This analysis revealed 628 and 1,113 differentially expressed genes (DEGs) in PI 88788 vs. Williams 82 and Forrest vs. Williams 82 comparisons, respectively (Figure 4A). Distinct transcriptional dynamics were observed between the two resistant cultivars. In PI 88788, approximately 65.9% of DEGs showed sustained upregulation following infection, suggesting a prolonged and potentially adaptive transcriptional response (Figure 4B). Gene Ontology (GO) enrichment analysis linked these DEGs to term such as “defense response, incompatible interaction”, “sucrose transport”, and “induced systemic resistance, ethylene mediated signaling pathway”. In contrast, 80.6% of DEGs in Forrest were sharply upregulated at 3 dpi, with enrichment in “cellular oxidant detoxification”, “response to defenses of other organism”, and “response to endoplasmic reticulum stress”, indicating an acute, immediate transcriptional reaction (Figure 4C). Canonical resistance genes, including those at *Rhg1* locus and *GmPR10-09g*, showed significantly elevated expression in both resistant cultivars compared to Williams 82 at one or more time points (Figure 2D–G). By contrast, other resistance-associated genes such as *Glyma.08G108900*, *Glyma.02G054200*, and *Glyma.01G147600* were upregulated in the xylem, cortex, and pericycle of resistant cultivars (Figure S4), but not in syncytial cell, likely due to their low expression in these cells (Figure 3D, E, I).

**Figure 4.**
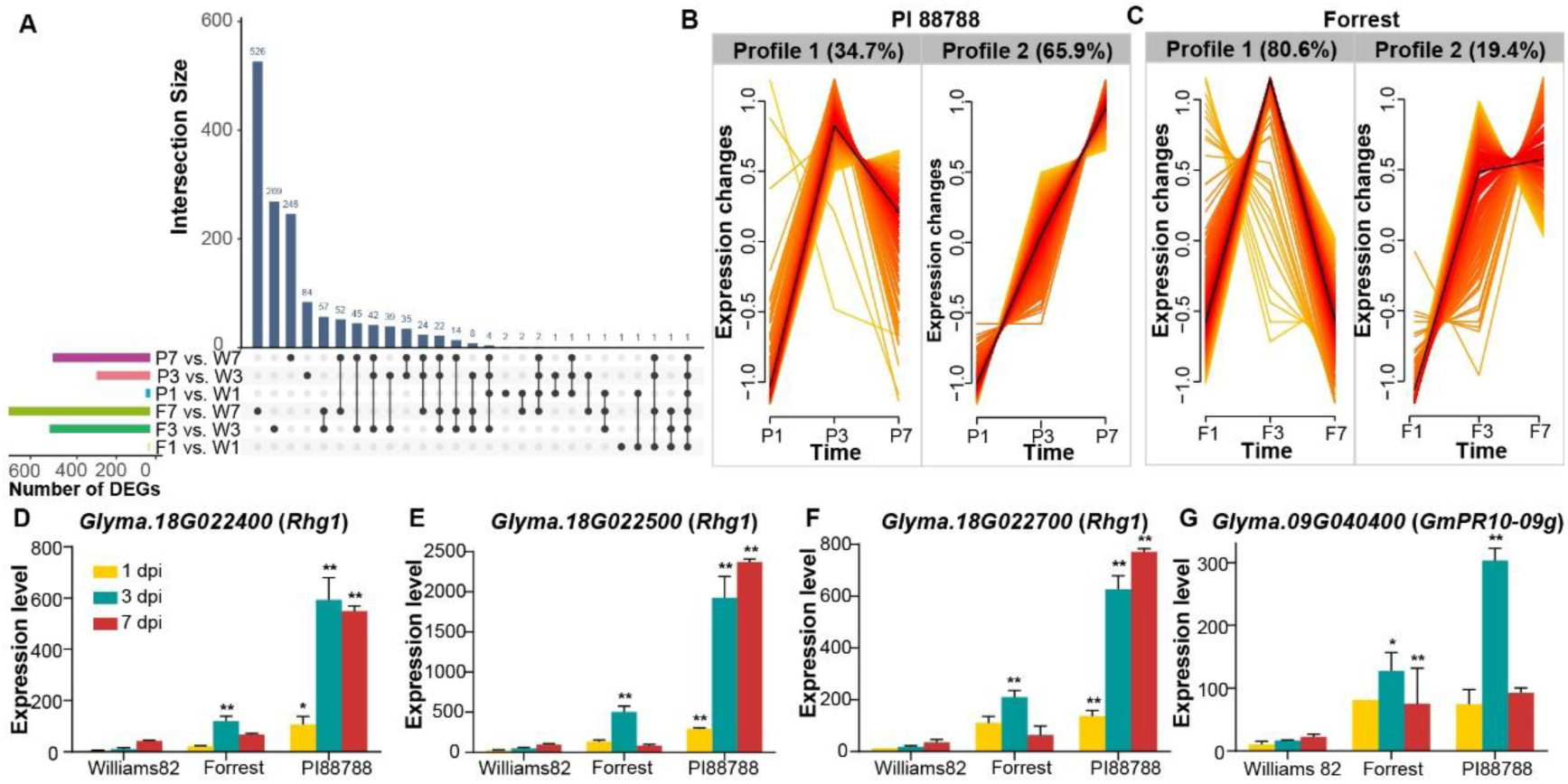
Comparative expression analysis of SCN-responsive genes among soybean cultivars. (A) Significantly differentially expressed genes (DEGs) identified between resistant and susceptible cultivars at three time points post inoculation with SCN. (B) Gene expression profiles of DEGs between Forrest and Williams 82. The number of genes in each profile is indicated above each box. F1: Forrest at 1 day post inoculation (dpi); F3: Forrest at 3 dpi; F7: Forrest at 7 dpi; P1: PI 88788 at 1 dpi; P3: PI 88788 at 3 dpi; P7: PI 88788 at 7 dpi. (C) Gene-expression profiles of DEGs between PI 88788 and Williams 82. The number of genes in each profile is indicated above each box. W1: Williams 82 at 1 day post inoculation (dpi); W3: Williams 82 at 3 dpi; W7: Williams 82 at 7 dpi. (D) –(G) Histograms showing the expression patterns of four functional genes reported by previous studies in the syncytium. * indicated that the *P*-value calculated using DESeq2 was less than 0.05, and ** indicated that the *P*-value was less than 0.01.

We further identified 383 novel syncytium-specific genes that were significantly upregulated in resistant cultivars (Table S5, Figure 5A–C). Functional validation *via* hairy root transformation assays confirmed three candidates. Overexpression of *Glyma.04G100400* (a phosphate-responsive 1 family gene), *Glyma.10G212900* (a GTP-binding elongation factor Tu family gene, *GmEF9*) (Gao et al., 2019), and *Glyma.15G013900* (a coat protein complex-related gene) significantly delayed nematode development in transgenic hairy roots of Williams 82 (Figure 5D, E). To explore the broader regulatory context, weighted gene co-expression network analysis (WGCNA) identified 19 gene modules in Forrest at 3 dpi (Figure S5A). The turquoise module, encompassing the three validated genes and 1,921 others, was enriched in processes such as “ATP metabolism”, “glucose metabolism”, “cell wall biogenesis”, and “nucleotide phosphorylation” (Figure S5B). Interestingly, using each of these three functional genes as hub node, we found top 10 strongly connected genes with them were coincident, highlighting potential regulatory partners involved in syncytium-mediated resistance. Therefore, a co-expression network including three functional genes and 10 coincident genes were constructed (Figure 5F). Notably, *Glyma.11G013200* was hypothesized to respond to SCN (Qu et al., 2025).

**Figure 5.**
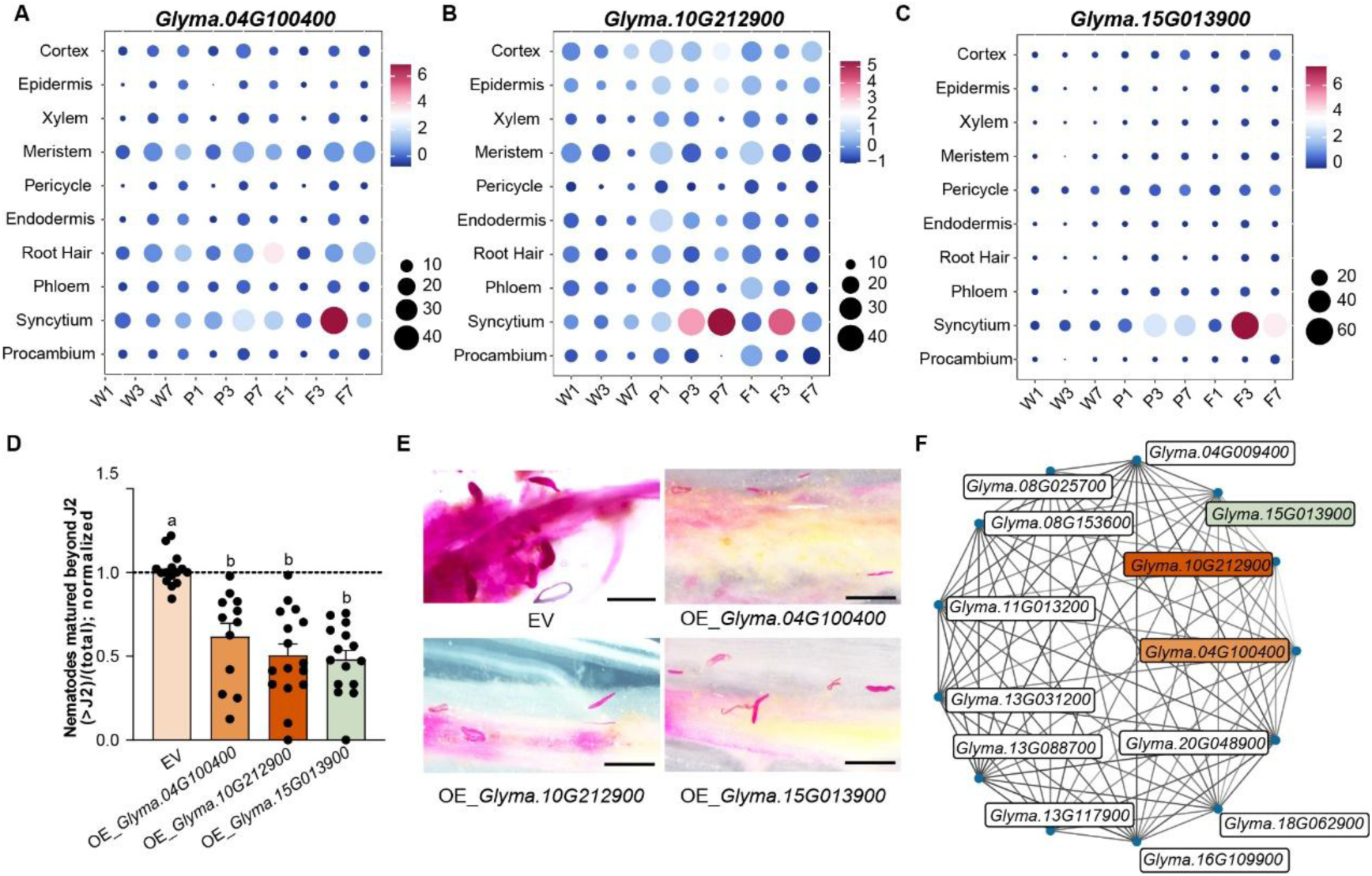
Overexpression of three novel candidate genes significantly suppressed SCN development. **(A) –(C)** Dot plot showing expression patterns of *Glyma.04G100400*, *Glyma.10G212900*, and *Glyma.15G013900* in syncytial cells in each sample, respectively. **(D)** Boxplot showing the ratio of nematodes matured beyond the J2 stage (J3 + J4) to total nematode number in transgenic hairy roots with empty vector, *Glyma.04G100400, Glyma.10G212900, and Glyma.15G013900* overexpression, respectively. The number of nematodes at the J2, J3, and J4 developmental stages was counted at 14 dpi, and the data are displayed as box-whisker plots with individual data points. Ratios in the EV were normalized to 1. Groups labeled with different letters are significantly different (*P* < 0.01, one-way ANOVA). **(E)** Overexpression of *Glyma.04G100400*, *Glyma.10G212900*, and *Glyma.15G013900* enhanced nematode resistance in transgenic hairy roots. Roots were stained with acid fuchsin. Bar = 500 µm. The above experiments were performed 3 times with similar results. (F) The co-expression networks of three functional genes and its co-expression genes. The depth of the lines represents the strength of the co-expression relationships.

### Reconstruction of the re-differentiated trajectory from procambial cells to initial syncytial cells

Syncytial cells were detected in all cultivars at 1 dpi, confirming the early induction of initial syncytial cells (ISCs). To identify the progenitor cell type undergoing re-differentiates into the syncytia, we extracted syncytial cells along with vascular bundle-associated cells, including endodermis, pericycle, procambium, phloem, and xylem for further comparative analysis. Pearson correlation analysis revealed that procambial cells had the highest transcriptional similarity to syncytial cells at 1 dpi (Figure 6A), suggesting that procambium cells serve as the development origin of syncytia.

**Figure 6.**
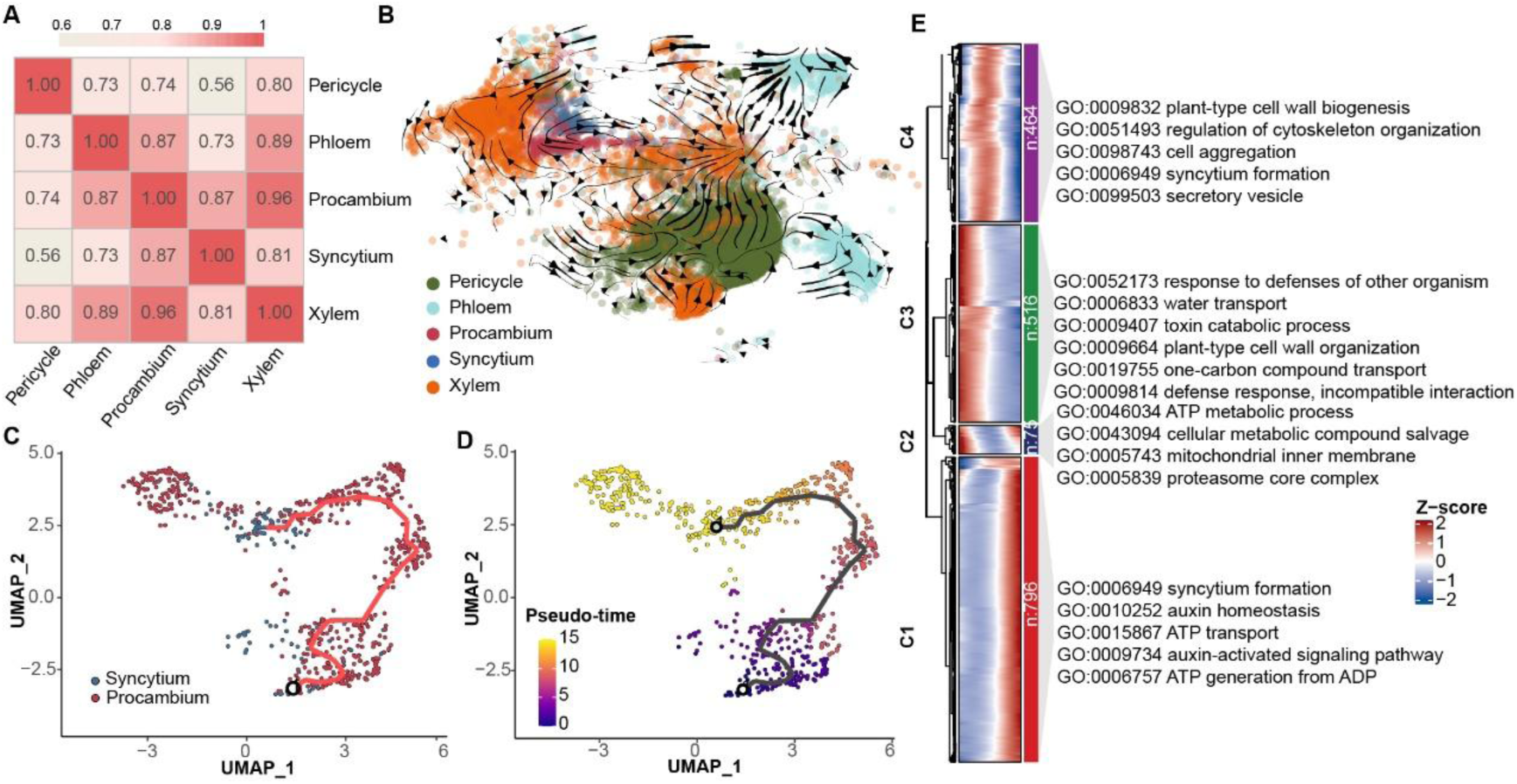
The re-differentiated process from procambial cell to syncytial cell. (A) Heatmap depicting gene expression correlation among pericycle, phloem, procambium, syncytium, and xylem at 1 day post-inoculation with soybean cyst nematode. (B) Developmental trajectories from procambial cells to syncytial cells inferred by scVelo using a dynamical model. Arrows indicate differentiation streams. (C), (D). Pseudo-time trajectories of the re-differentiation process from procambial cells to syncytial cells. Cells are colored by cell type (C) or developmental pseudo-time (D), respectively. (E). Expression heatmap of differentially expressed genes along the pseudo-time trajectory. Enriched Gene Ontology (GO) terms are shown in the right panel.

To reconstruct the developmental trajectory, we applied scVelo (Bergen et al., 2020), which delineated a clear transition pathway from procambium cells to syncytial cells (Figure 6B; Figure S6A). Pseudotime analysis using Monocle3 (Cao et al., 2019) further supported this trajectory, identifying two subclusters (PS1, PS2): PS1 comprised a mix of procambial and the putative ISCs, whereas PS2 was enriched for syncytial cells (Figure S6B, C). Using procambial cells in PS1 as the trajectory root, we identified 1,851 genes associated with the re-differentiation processes (Figure 6 C, D). Early in pseudotime analysis, upregulated genes were enriched in pathways related to “response to organisms”, “defense response (incompatible interaction)”, and “one-carbon transport”, indicating initial defense response during SCN penetration of the vascular. As differentiation progressed, genes associated with “syncytium formation”, “cell wall organization”, and “cell aggregation” were upregulated, consistent with known cellular remodeling events during syncytium establishment (Golinowski et al., 1996; Sobczak *et al*., 1997b; a). Concurrently, genes involved in ATP transport and metabolism were strongly induced, highlighting the increased energy demand in developing syncytia. Additionally, auxin signaling pathways were prominently activated, underscoring their role in promoting syncytial reprogramming and differentiation (Figure 6E; Table S6). Collectively, these data provide robust molecular evidence, complemented by classical histological and ultrastructural studies, that procambial cells are the primary origin of syncytial cells during early SCN infection. This integrated approach not only reinforces previous morphological observations but also advances our mechanistic understanding of the early cellular transitions underlying syncytium formation.

### Divergent developmental trajectories and heterogeneity of syncytial cells across soybean cultivars

At 7 dpi, syncytial cell counts were markedly lower in Forrest compared to PI 88788 and Williams 82. This reduction likely reflects the early arrest of syncytium development in Forrest, where 67.74% of syncytia were terminated by 3 dpi (Figure S7A). This observation aligns with morphological studies showing that *rhg1-a* and *Rhg4*-mediated resistance in the Peking-type response is activated rapidly, within 48 hours of infection, while *rhg1*-b mediated resistance in PI 209322 (the source of PI 88788-type resistance) initiates later, around 8 to 10 dpi (Kandoth et al., 2011; Mahalingam and Skorupska, 1996). Following their initial formation, syncytia expand by incorporating adjacent host cells. Re-clustering of syncytial cells revealed three subclusters (Syn0, Syn1, and Syn2), reflecting transcriptional heterogeneity (Figure S8B). UMAP visualization confirmed that this heterogeneity was driven by both cultivar and infection stage (Figure S7C–E).

To dissect these developmental dynamics, we reconstructed cultivar-specific syncytial cell trajectories using Monocle3 (Figure 7A–F). All trajectories followed a progression from 1 to 3 to 7 dpi, yet the underlying gene expression patterns differed substantially, particularly in Forrest. In both Williams 82 and PI 88788, genes associated with defense response and gene silencing were upregulated early in the trajectory. No major transcriptomic changes were observed at intermediate stages. By contrast, genes involved in energy metabolism, including “ATP metabolic and biosynthetic processes” and “glycolytic processes”, were activated at later stages (Figure 7G, H). In Forrest, defense-related genes were also induced early, along with those involved in syncytium formation, ATP production, and fluid transport. Notably, more than 50% of pseudotime-dependent genes peaked during intermediate stages, with enrichment in pathways related to ethylene and jasmonic acid signaling, vesicle-mediated transport, and regulation of programmed cell death. This pattern, coupled with the early arrest of syncytia (Figure S7A), suggest that these processes contributed to syncytium degeneration in Forrest. At later stages, energy metabolism genes, particularly those linked to “ATP metabolic and biosynthetic processes” and “glycolytic processes” were again strongly expressed (Figure 7I), indicating high metabolic activity in mature syncytial cells.

**Figure 7.**
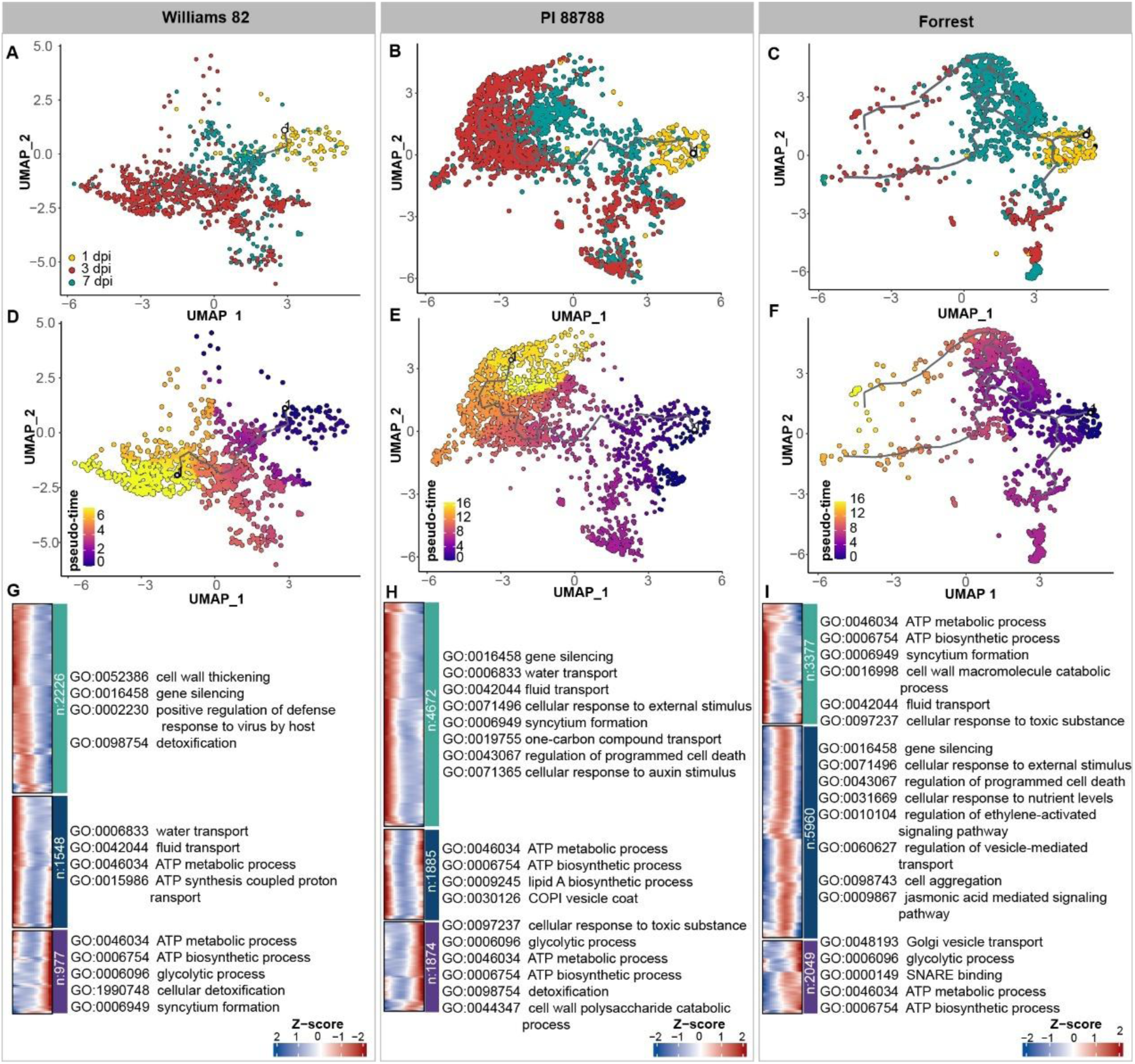
Developmental trajectories of syncytial cells in Williams 82, PI 88788 and Forrest, respectively. (A)–(F). Pseudo-time trajectories of syncytial cells inferred by Monocle 3 in Williams 82 (A, D), PI 88788 (B, E), and Forrest (C, F), respectively. Cells are colored by sample time (A–C) or pseudo-time (d–f). **(G)–(I**). Expression heatmaps showing expression patterns of pseudo-time-dependent genes along the pseudo-time trajectory in Williams 82 (G), PI 88788 (H), and Forrest (I), respectively. Enriched Gene Ontology (GO) terms are shown in the right panel.

## Discussion

In this study, we performed snRNA-seq on 172,199 high-quality nuclei from three soybean cultivars across three SCN infection stages (Figure 1A), generating the first single-cell transcriptional atlas of soybean root response to SCN. Through integrative pseudotime trajectory analyses, we reconstructed the re-differentiation pathway from procambial cells to initial syncytial cells, providing insight into the cellular origin and developmental progression of syncytia across resistant and susceptible cultivars.

Our custom dual-species genome mapping strategy, which incorporated both soybean and SCN genomes, enabled us to discriminate between host- and nematod-derived nuclei. Although fewer SCN-derived nuclei were recovered than expected (Table S1; Figure S1C), likely due to protocol incompatibilities, the high proportions of soybean nuclei captured validated the robustness of our plant-focused extraction and processing workflow. These findings highlight the need for optimized protocols to improve the recovery of nematode nuclei in future dual-species single cell studies, which will enhance our understanding of the precious mechanism governing soybean-nematode interactions at single cell level.

By integrating four complementary annotation strategies, we identified ten distinct root cell types, including syncytia (Figure 2A–E). This comprehensive annotation enabled the spatial mapping of gene expression and identification of syncytium-enriched genes associated with SCN resistance. Consistent with their functions as feeding sites, syncytial cells showed enrichment for genes involved in ATP biosynthesis, metabolic activity, and hormone pathways (Figure 2F), underscore their role in nutrient transfer to the nematode.

A major advance of this work is the integration of spatial and temporal transcriptomics, which enabled high-resolution dissection of SCN resistance gene expression. Known resistance genes exhibited distinct spatial expression across multiple root tissues (Figure 3; Figure S3). For example, *GmSNAP18* was significantly enriched in the syncytia of PI 88788 and Forrest, consistent with *in situ* hybridization (Lakhssassi *et al*., 2020) and immunolabeling studies (Bayless *et al*., 2016; Bayless *et al*., 2019) (Figure 3A). Two additional *Rhg1* genes, *Glyma.18G022400* and *Glyma.18G022700*, displayed enrichment in the epidermis and endodermis, respectively, of PI 88788, supporting their roles at the early infection of SCN (Figure 3B, C).

Unlike *Rhg1*, *Rhg4* gene (*GmSHMT08*) showed no significant differential expression in syncytia between resistant and susceptible cultivars (Figure 3D), consistent with previous reports that its resistance function is conferred through protein-coding SNPs rather than transcriptional induction (Patil et al., 2019). It exhibited elevated expression in xylem of resistant cultivars and in the syncytia of Forrest at 1 dpi, suggesting a role in vascular defense (Figure S4A). Similarly, *GmPR10-09g* was highly expressed in syncytia and cortex of resistant cultivars at 3 dpi, validating both our annotation and the multi-cellular nature of SCN resistance (Figure 3F).

Through comparative analysis of syncytium-specific DEGs between resistant and susceptible cultivars, we identified 383 candidate resistance genes uniquely enriched in syncytial cells (Table S5). Several candidates, such as *Glyma.08G019300* (reticulon-like protein 2). and *Glyma.13G240100*/*Glyma.15G073200* (lipid droplet-associated proteins), are previously known to be involved in biotic and abiotic stress responses (Gidda et al., 2016; Lee et al., 2011). Notably, three novel genes-*Glyma.04G100400*, *Glyma.10G212900* (*GmEF9* (*Gao et al., 2019*)), and *Glyma.15G013900*-displayed syncytium-enriched expression specifically in the resistant cultivars Forrest and PI 88788. Overexpression of these genes in transgenic hairy roots significantly impaired nematode development (Figure 5). Intriguingly, these genes have been previously implicated in diverse stress-related pathways, including drought, salinity, ABA signaling, and pathogen defense (Fu et al., 2012; Li et al., 2018; Mendez-Lopez et al., 2023; Wu et al., 2018; Zhang et al., 2022), suggesting that SCN resistance mechanisms may co-opt broad-spectrum stress response networks. There were ten *EF1α* family members have been annotated in the soybean genomes, among which *GmEF9* has been previously characterized for its role in stress adaption (Gao *et al*., 2019). Pseudotime analyses further revealed that these genes peaked during later syncytial development stages in resistant cultivars but not in susceptible cultivar Williams 82 (Figure 7), reinforcing their potential involvement in cultivar-specific active defense mechanisms. Initial syncytial cells were able to formatted in both resistant and susceptible soybean cultivars. By integrating scVelo and Monocle3, we reconstructed the re-differentiated trajectory from procambial cells to initial syncytial cells, identifying 1,851 re-differentiation-associated genes enriched in early defense, cell wall reorganization, metabolic activity, and auxin signaling (Figure 6; Figure S6). This is consistent with classical histological studies in *Arabidopsis*, in which juvenile *Heterodera schachtii* females were observed to target and penetrate procambial cells to initiate syncytium formation (Golinowski *et al*., 1996; Sobczak *et al*., 1997b). These studies reported ultrastructural changes, such as alterations to the cell wall and vacuolar organization, that mirror the transcriptional shifts identified in our dataset, indicating early cellular reprogramming during syncytium initiation.

After the establishment of ISC, distinct transcriptional responses accrued in Forest and PI 88788 (Figure 4B, C). In Forrest, a rapid incompatible reaction to SCN resulted in syncytium arrest by 3 dpi, whereas PI 88788 displayed prolonged syncytial development through 7 dpi (Figure 7; Figure S7A). Genes associated with ethylene and jasmonic acid signaling, vesicle transport, and programmed cell death were uniquely upregulated at intermediate stages in Forrest, implicating their roles in syncytium degeneration. Given the observation that syncytia undergo degeneration by 10 dpi (Kandoth *et al*., 2011), we hypothesize that snRNA-seq datasets of PI 88788 collected around the 10 dpi will facilitate a more comprehensive understanding of the molecular mechanisms underlying syncytium degeneration.

Taken together, our study presents a high-resolution single-cell transcriptional atlas of soybean root responses to SCN infection. We identified novel syncytium-specific resistance genes, traced the re-differentiation of procambial cells into syncytia, and revealed cultivar-specific trajectories associated with syncytial development and arrest. These insights advance our understanding of host-pathogen interactions and provide a valuable resource for soybean breeding programs targeting enhanced SCN resistance.

## Methods

### Plant materials and nematode inoculation

The seeds of soybean cultivars Williams 82, PI 88788, and Forrest were sterilized by soaking in 10% sodium hypochlorite for 10 minutes. Subsequently, they were planted in high-temperature sterilized vermiculite at 26°C under a 14:10 light-dark cycle. After 5 days, seedlings with consistent growth conditions were gently removed from the hypochlorite solution and placed on moistened germination papers. SCN eggs of HG-type 0 (race 3) were incubated in sterile water at 28°C for 3 days. Approximately 400 infective J2 SCN were applied 1 cm above the root tip. After 24 hours of inoculation, the germination papers were rolled into rag dolls and placed in the plant growth chamber at 26°C with a 14:10 light-dark cycle. The infection process of SCN was examined using 20% of the inoculated samples using acid fuchsin stain.

### Nuclei isolation, 10×snRNA-seq library construction, and snRNA sequencing

For single-nucleus RNA sequencing (snRNA-seq), the infection zone of the root with obvious infection trace at 1, 3, and 7 days post-inoculation (dpi) was collected, with two biological replicates per condition. The chopped root was added to a centrifuge tube containing ice-cold 1× Nuclei Isolation Buffer (NIB), and the sample was centrifuged at 300 × g for 1 minute. The supernatant was then transferred to a new centrifuge tube, and the isolated nuclei were sorted using fluorescence-activated nucleus sorting (FANS). After assessing the nuclei quality and counting them under a microscope using the DAPI channel, > 16,000 nuclei per sample were loaded onto the 10x Genomics chip. Sequencing was conducted on the Illumina Novaseq6000 platform in PE150 mode.

### Processing of snRNA-seq raw data

The procession of snRNA-seq data was performed using Cell Ranger (version 7.2.0) using the default parameters (Zheng et al., 2017). A combined reference genome of soybean genome (Wm82.v4; download from https://www.soybase.org/) and SCN genome (ASM414822v2) was manually created using the “cellranger mkref” command. Unique Molecular Identifiers (UMIs) were quantified based on the manually combined soybean GTF file and SCN GTF file using “cellranger count.”

### Data integration, and clustering

Cell-by-gene matrices were subsequently imported into the R package Seurat (version 5.0.3) (Hao *et al*., 2024) for the following analysis. For quality control, nuclei and genes were filtered based on the following criteria: (1) according to the distribution of genes expressed, genes that expressed in fewer than five nuclei were filtered out; (2) according to the distribution of unique gene counts in each nuclei, we filtered nuclei with unique gene counts between 450 and 15,000 of Forrest at 7 dpi and Williams 82 at 7 dpi; and for the other samples, nuclei with unique gene counts between 450 and 100,000, and UMI counts below 25,000 were retained; (3) DoubletFinder v2.0.4 (McGinnis et al., 2019) was used to identify the potential doublets according to its manual. (4) a nucleus was classified as soybean- or SCN-derived if > 80% of its expressed genes were from soybean or SCN, respectively, and the remaining nuclei were filtered.

Data normalization was performed using the “NormalizeData” function. The top 2,000 highly variable genes identified by “FindVariableFeatures” were used for principal component analysis (PCA) dimensionality reduction. All samples were integrated using the “IntegrateLayers” function in Harmony (Korsunsky et al., 2019). The first 30 principal components (PCs) were used to construct clusters with a resolution of 1. The Uniform Manifold Approximation and Projection (UMAP) method was applied for cluster visualization. Syncytial cells were extracted, and the first 20 PCs were re-clustered with a resolution of 0.1.

### Cell type annotation

Differentially expressed genes for each cluster were identified using the Wilcoxon rank-sum test via the “FindAllMarkers” function in Seurat (version 5.0.3). Parameters were set as follows: (1) genes expressed in > 25% of the nuclei in a cluster; (2) avg_log2FC of up-regulated genes ≥ 0.5. The biomarkers for each cluster were compared with verified markers from soybean and *Arabidopsis thaliana* in the literature (Table S2, Table S3). *Arabidopsis* homologs of the biomarkers were uploaded into the scPlantDB database to predict the cell type using the “Cell Type Predictor” tool (https://biobigdata.nju.edu.cn/scplantdb/home) (He *et al*., 2024).

For the Seurat-based cell type annotation, a processed soybean root dataset was downloaded from the GEO (Accession number: GSE270392). The data were re-normalized using Seurat’s “NormalizeData” function. After mapping our dataset to the reference dataset using the “FindTransferAnchors” function, cell types were predicted using the “TransferData” function. To account for cell-type imbalances, the prediction results were normalized by dividing the total number of cells predicted for each cell type by the number of cells predicted within each cluster.

For the cell type annotated using Garnett (version 0.1.23) (Pliner *et al*., 2019), a custom classifier was trained using known cell-type-specific marker genes listed in Table S2. After revising the marker file based on ambiguity scores using the “check_markers” function, the classifier was trained with the “train_cell_classifier” function. Finally, the “classify_cells” function was used to classify the cells, and the prediction results were normalized in a similar manner.

The cell type-specifically expressed genes were identified using Cellex (Timshel et al., 2020), with only those exhibiting a specificity score > 0.6 being retained.

### Differential gene expression analysis

DESeq2 (version 1.42.1) (Love et al., 2014) was utilized to perform the pseudo-bulk differential expression analysis according to the tutorial from the Harvard Chan Bioinformatics Core (https://hbctraining.github.io/scRNA-seq/lessons/pseudobulk_DESeq2_scrnaseq.html). For the syncytial cells, the sample-level meta-data and count in the resistant and susceptible cultivars collected on the same day were aggregated to run the DESeq2. i.e., F1 vs. W1, F3 vs. W3, F7 vs. W7, P1 vs. W1, P3 vs. W3, P7 vs. W7. Genes with *P*.adj < 0.05 and |log2FoldChange| > 1 were considered significantly differentially expressed. After that, the SCT normalized expression of DEGs between Forrest and Williams 82, as well as PI 88788 and Williams 82 were used to do the time series expression pattern analysis using R package Mfuzz (version 2.64.0) (Kumar and M, 2007).

### Correlation analysis

The correlation analyses for the transcriptome of each sample and the transcriptome of each cluster in the 1 dpi were performed using the R function “cor”. We used Euclidian to calculate the distance, and the ‘‘pearson” cluster method to compute the correlation coefficient.

### Sing-cell trajectory analysis

The trajectory of transition from the procambium cells to the syncytial cells was constructed using the Monocle 3 (version 1.3.7) (Cao *et al*., 2019). The Seurat object of the two cell types in the 1 dpi was first converted into the Monocle 3 using the “new_cell_data_set” function. The “preprocess_cds”, “align_cds”, “reduce_dimension”, and “cluster_cells” functions were used for standardization, reduction, and clustering of the Monocle 3 object. The trajectory inference was performed using the “learn-graph” function and visualized using the “plot-cells” function. Differentially expressed genes along the trajectory were clustered and visualized using ClusterGVis (https://github.com/junjunlab/ClusterGVis).

### RNA velocity analysis

The RNA velocity was performed using the Python package scVelo (version 0.2.4) on the dynamical model (Bergen *et al*., 2020). The spliced and unspliced matrices were first calculated using velocyto (version 0.17.17) (La Manno et al., 2018). After that, the count, metadata, embedding information of UMAP, and gene name in the Seurat object were converted to the scanpy object, i.e., adata (Wolf et al., 2018). Then, RNA velocity analysis was carried out according to the provided tutorials.

### Gene enrichment analysis

For gene function annotation, the protein sequences of the soybean genome (Wm82.v4) were mapped against the EGGNOG 5 using eggNOG-mapper v2 (Cantalapiedra et al., 2021). The parameters were set as default, except for the Taxonomic Scope was specified as the Viridiplantae. Then, Gene ontology enrichment analysis was performed using clusterProfiler v4.10.1 (Wu et al., 2021) with a *P*-value cutoff of 0.05.

### Weighted-correlation network analysis

The weighted-correlation network analysis was performed using the hdWGCNA v0.4.06 following the manual (Morabito et al., 2023). Give the significant expression levels of three novel functional genes in the Forrest at 3 dpi, subset of syncytial cells was extracted from this sample. Genes expressed in at least 5% of cells were selected using the “SetupForWGCNA” function, serving as the basis for subsequent analyses. Metacells were then generated through the “MetacellsByGroups” function, with the parameter k set to 25. The soft-power threshold was set as 4 according to the scale-free graph. Subsequently, the co-expression network was constructed using the “ConstructNetwork” function. Top 10 genes exhibiting the strongest connections with the target functional genes were identified based on the topological overlap matrix weights. The ggraph (Si et al., 2020) was employed to visualize co-expressed genes with functional genes.

### Gene cloning and vector construction

The overexpression vector was constructed following the protocol outlined by Chen et al., 2024, primers were firstly designed to target the 5’ UTR and 3’ UTR regions of the gene based on the Williams 82 cDNA reference. PCR was then performed using Forrest cDNA as the template (primers were listed in the Table S7). The PCR products were subsequently ligated into a T-vector, and 3-5 single bacterial colonies were selected for sequencing. Clones exhibiting high reproducibility and strong homology to the Williams 82 reference gene were further amplified and cloned into the pHS174 vector, which carries a CaMV 35S enhancer promoter and the tdTomato RFP reporter gene. The empty pHS174 vector served as the negative control (Chen et al., 2024).

### Generation of transgenic soybean composite hairy roots

The *Agrobacterium rhizogenes* strain K599 harboring the recombinant pHS174 construct was inoculated onto YEP solid medium supplemented with 50 μg/mL streptomycin sulfate and 100 μg/mL kanamycin, followed by a 48-hour incubation. Subsequently, bacterial cells were harvested and inoculated onto the wound sites of plant tissues. Seven-day-old Williams 82 and Forrest seedlings were decapitated 1 cm below the cotyledonary node using sterile surgical blades. Inoculated plants were maintained in vermiculite under high humidity at 28°C in a growth chamber. Primary hairy roots emerged within 15–20 dpi. The growth of hairy roots was subsequently verified using a fluorescence microscope (Olympus SZX16, Tokyo, Japan), identifying positive roots through RFP filters.

### Functional validation analysis

Positive transgenic soybean hairy roots were inoculated with about 400 infectious J2 SCN. At 14 dpi, nematode-infected root samples were stained with 0.1% acid fuchsin solution (w/v in lactic acid), and SCN at various developmental stages were counted under a stereomicroscope. The ratio of J3 + J4 to J2 + J3 + J4 was calculated. Data from transgenic and wild-type roots were statistically analyzed using the Wilcox test.

## Supplementary information

Document S1

Tables S1–S7

## Acknowledgments

We thank Dr. Zhijian Liu in Northeast Normal University for helpful suggestions on the manuscript. This work is supported by the Zhejiang Lab (Grant No. 2021PE0AC04), National Natural Science Foundation of China (32272478), and the Fundamental Research Funds for the Central Universities (226-2024-00220).

## Author contributions

X.F., L.W., and S.H. conceived and supervised the project. P.B., S.H., L.W., and X.F., wrote the manuscript with the contributions of all other co-authors. P.B. and L.C. collected the samples. P.B. handled snRNA-seq analyses. X.J., Y.H, and L.C. contributed to the analytical, molecular cloning and transformation work. All the authors read and approved the final manuscript.

## Supplemental Figures

**Figure S1.**
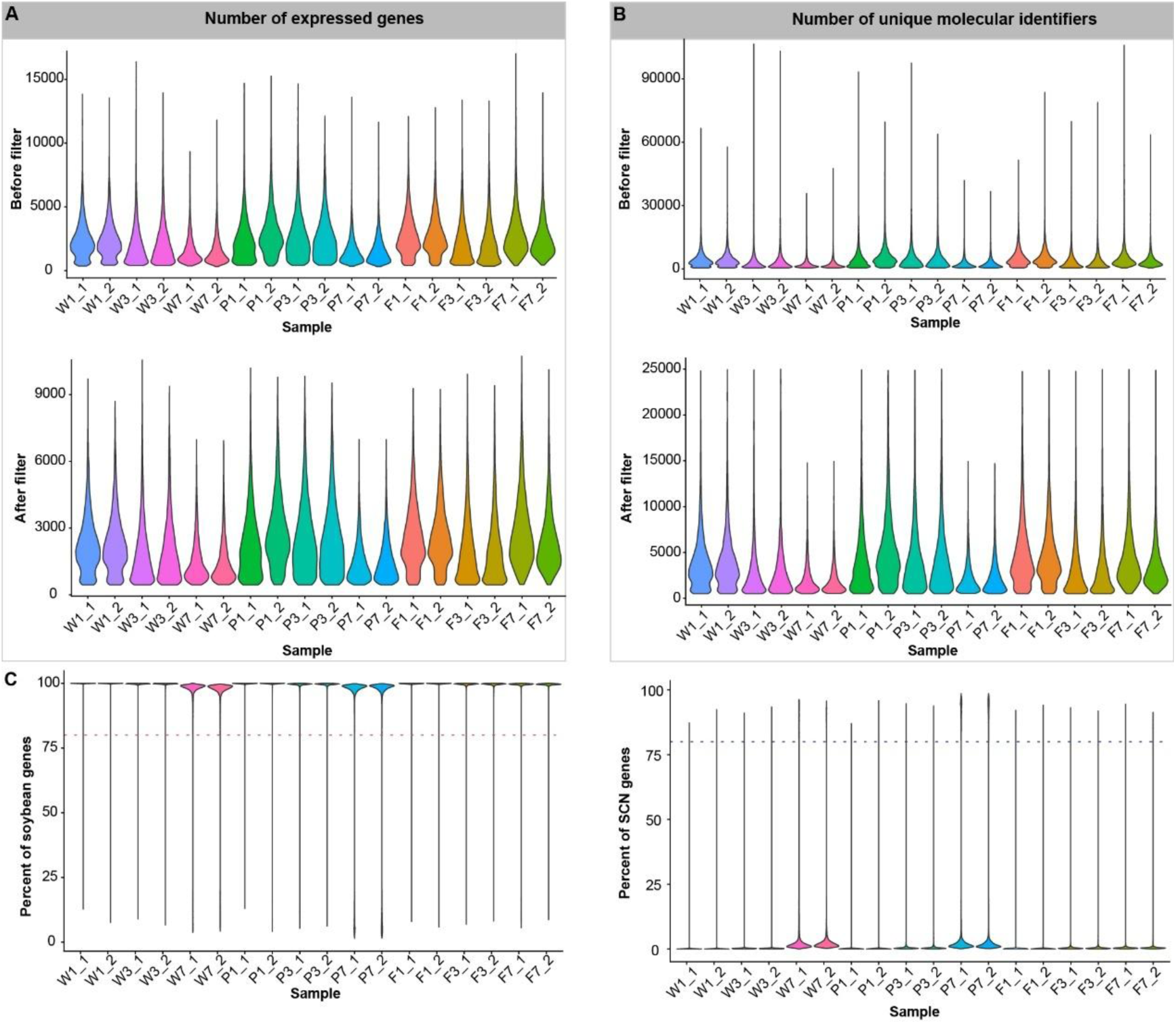
Quality control of snRNA-seq datasets. (A) Violin plots showing the distribution of gene counts per sample before and after quality control. (B) Violin plots showing the distribution of unique molecular identifiers (UMIs) per sample before and after quality control. (C) Violin plots showing the percentage of genes derived from soybean and SCN in each nucleus. F1: Forrest at 1 day post inoculation (dpi); F3: Forrest at 3 dpi; F7: Forrest at 7 dpi; P1: PI 88788 at 1 dpi; P3: PI 88788 at 3 dpi; P7: PI 88788 at 7 dpi; W1: Williams 82 at 1 dpi; W3: Williams 82 at 3 dpi; W7: Williams 82 at 7 dpi.

**Figure S2.**
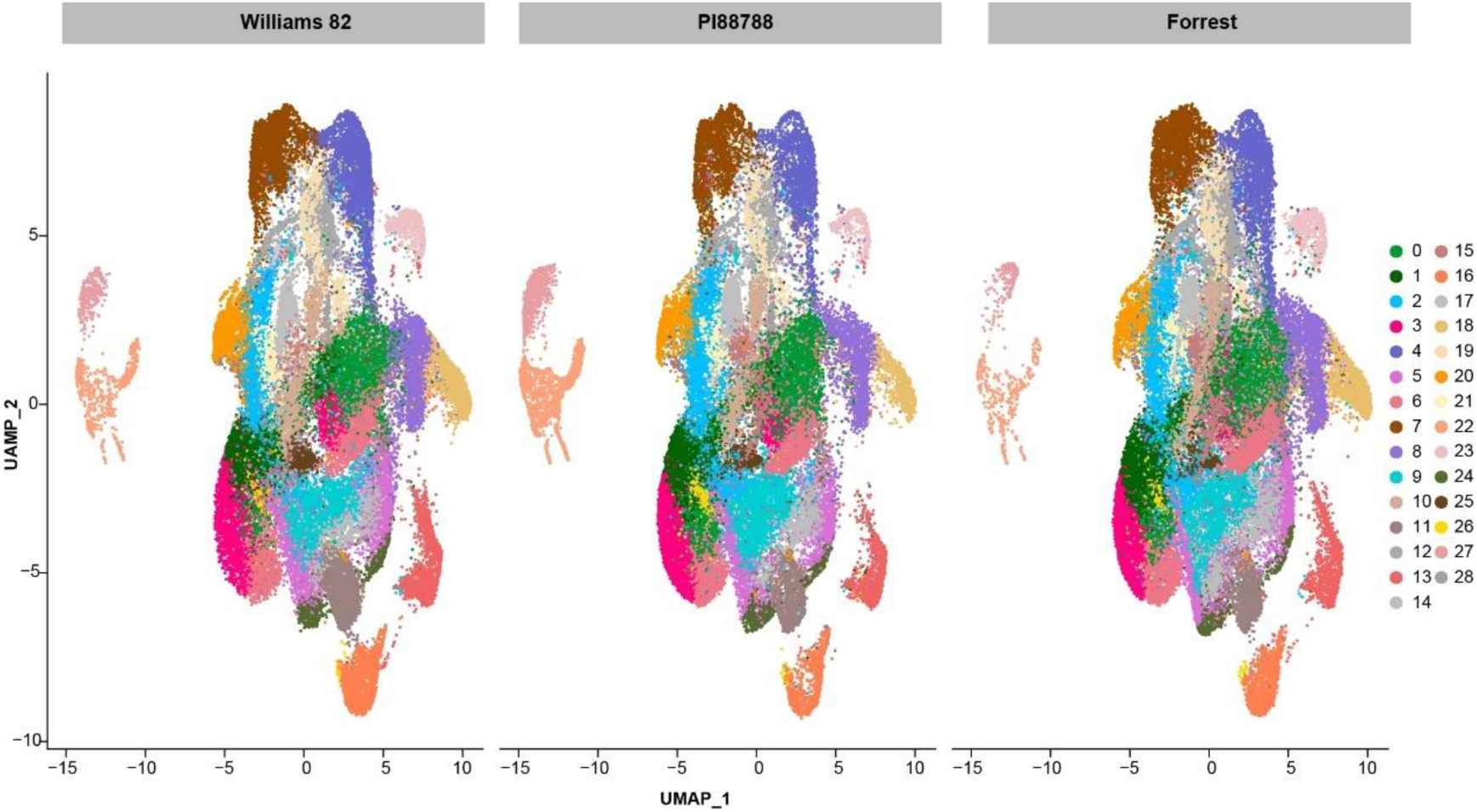
UMAP visualization of 29 clusters in soybean roots and soybean cyst nematode (SCN) from Forrest, PI 88788, and Williams 82, respectively.

**Figure S3.**
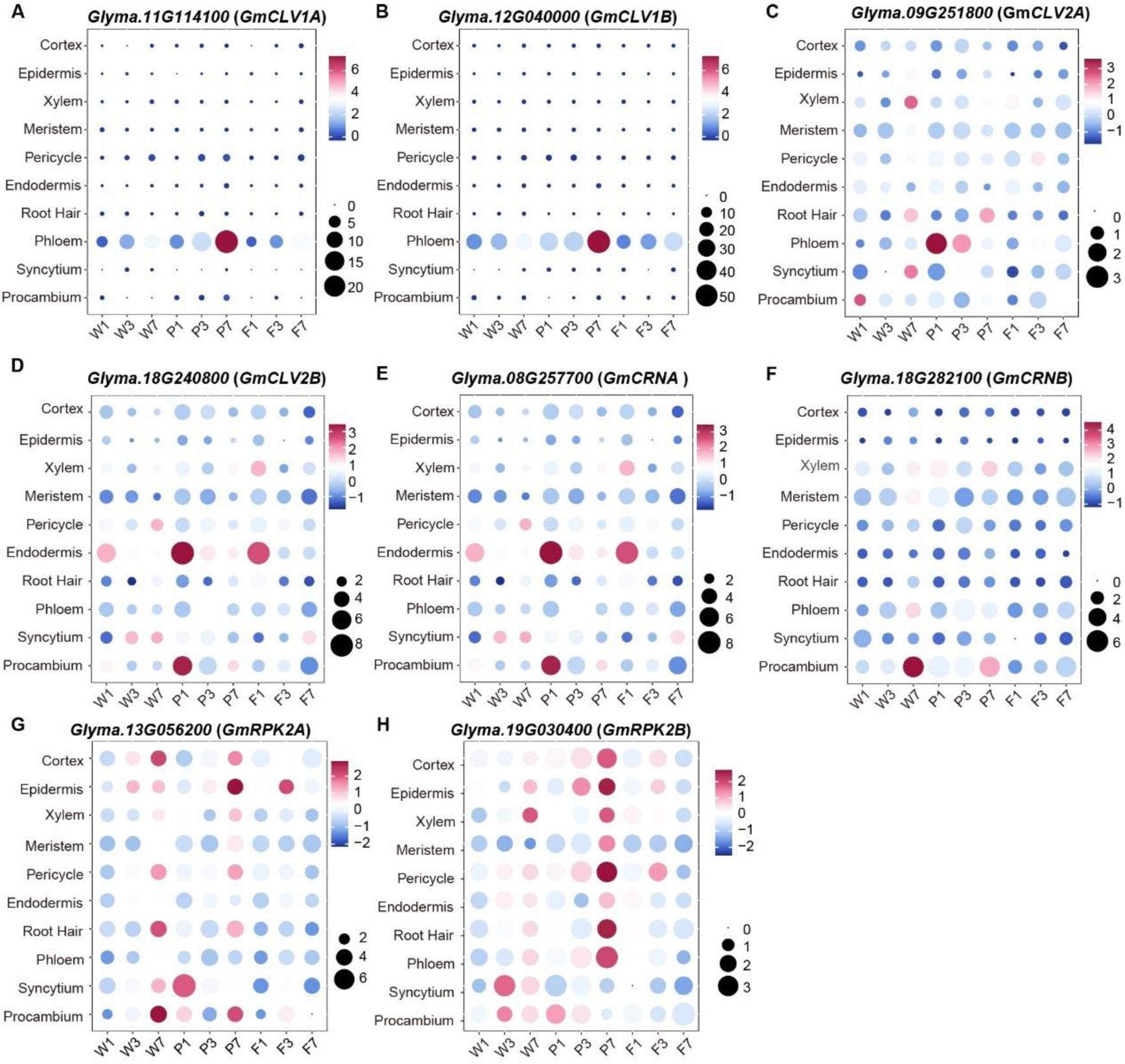
CLE receptors exhibited diverse root expression patterns.

**Figure S4.**
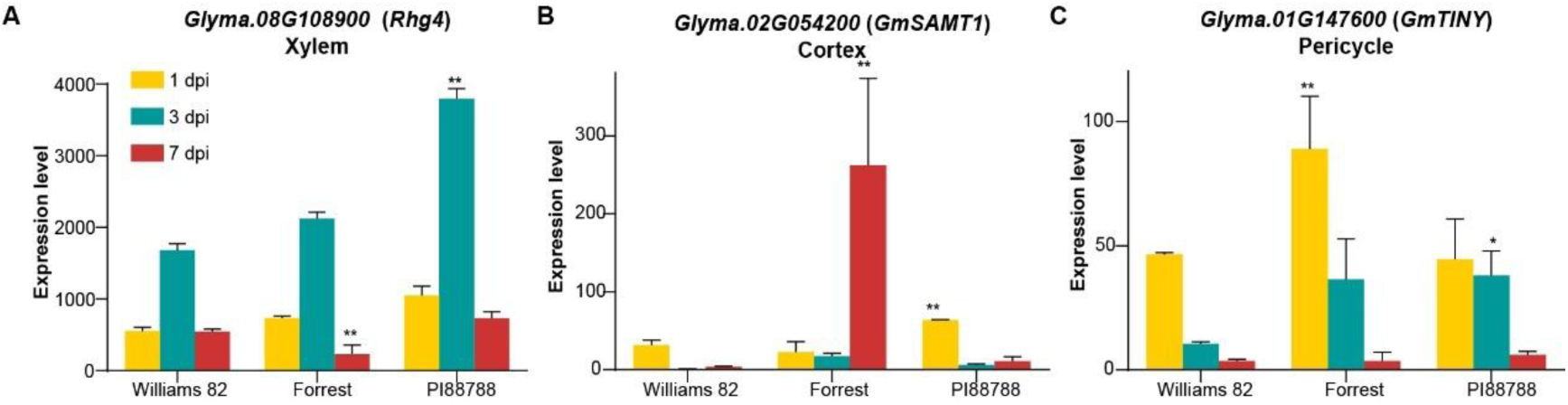
Expression profiles of three reported functional resistance genes. (A) Histograms depicting the expression profile of *Glyma.08G108900* in xylem. (B) Histograms depicting the expression profile of *Glyma.02G054200* in cortex. (C) Histograms depicting the expression profile of *Glyma.01G147600* in pericycle. Expression levels of these genes were extracted from cell types that they show the highest expression levels. * indicated that the *P*-value calculated using DESeq2 was less than 0.05, and ** indicated that the *P*-value was less than 0.01.

**Figure S5.**
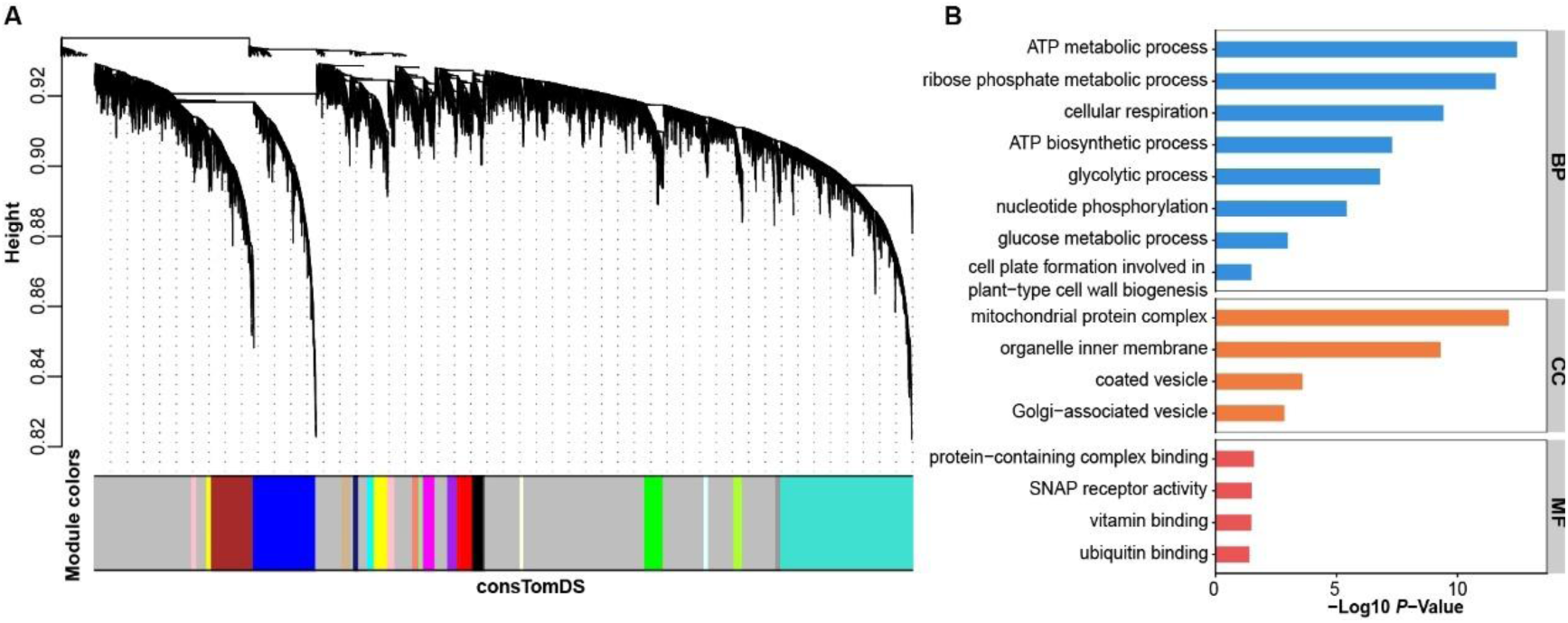
19 gene modules were found in Forrest at 3 days post inoculation. (A) Dendrogram plot showing the different co-expression modules resulting from the network analysis. The color at the bottom indicates the co-expression module assignment. (B) –(C) Partial gene ontology terms of genes in the turquoise module.

**Figure S6.**
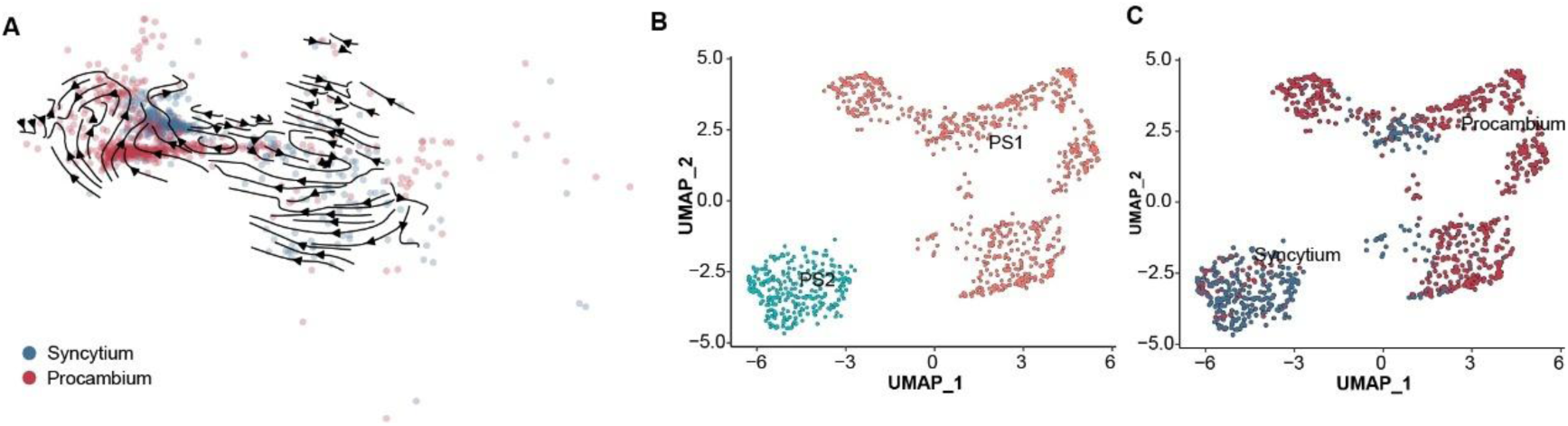
The re-differentiation process from procambial cells to syncytial cells. (A) Developmental trajectories from procambial cells to syncytial cells inferred by scVelo using the dynamical model. Arrows indicate differentiation streams. The arrows refer to the differentiation streams. **(B)** UMAP visualization of two subclusters: syncytial cell subclusters and procambial cell subclusters. Cells are colored by the subcluster. **(C)** UMAP visualization of two subclusters: syncytial cell subclusters and procambial cell subclusters. Cells are colored by the cell type.

**Figure S7.**
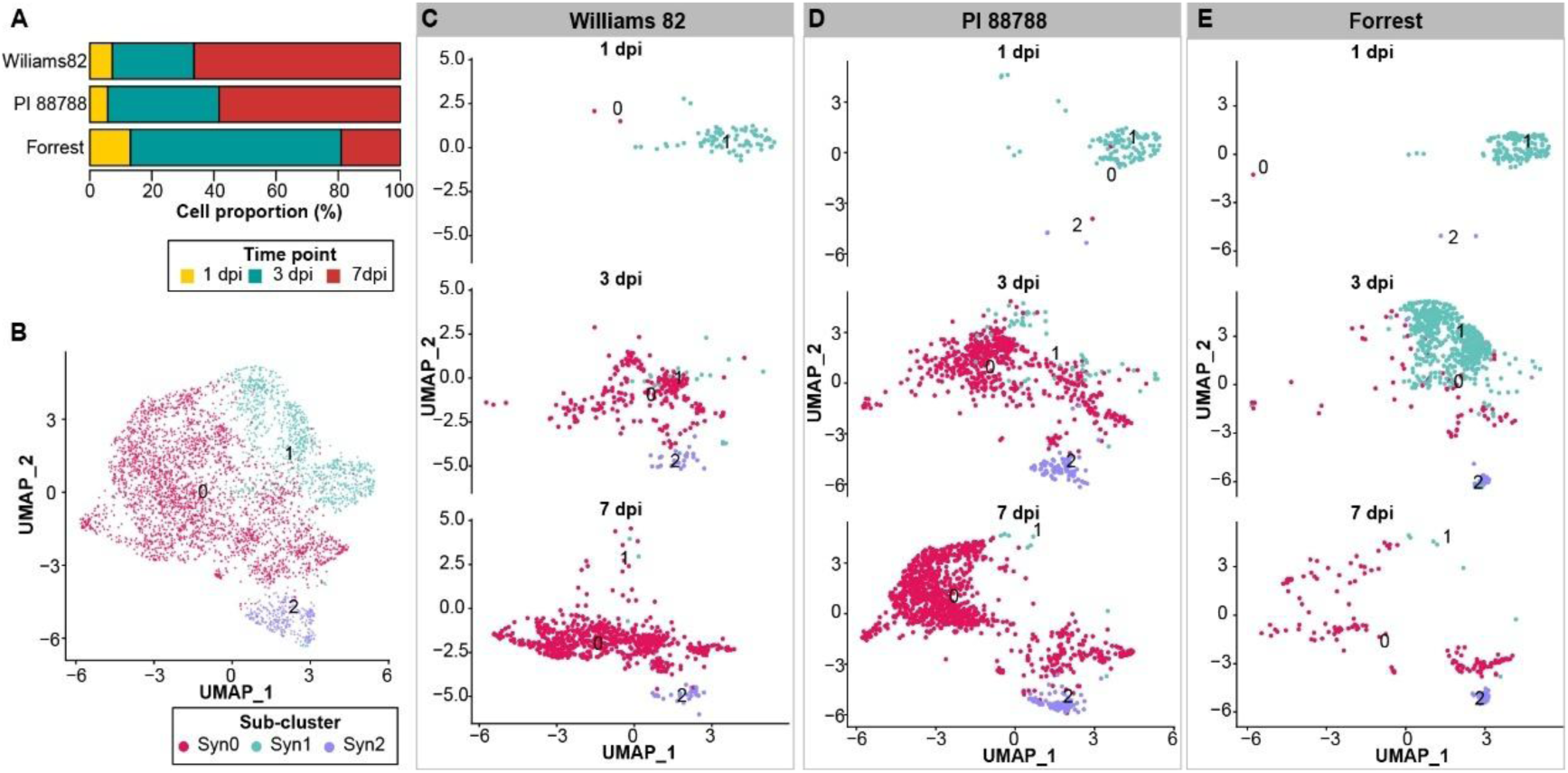
Syncytial cells demonstrated heterogeneity (A). Percentage of syncytial cells sampled at each time point across three cultivars. (B). UMAP visualization of three syncytial cell subclusters. (C)–(E). UMAP visualization of three syncytial cell subclusters at 1, 3 and 7 days post inoculation, respectively.

